# Subcellular Time Series Modeling of Heterogeneous Cell Protrusion

**DOI:** 10.1101/429118

**Authors:** Yeesock Kim, Hee June Choi, Kwonmoo Lee

## Abstract

In this paper, a new biological modeling approach is proposed for predicting complex heterogeneous subcellular behaviors. Cell protrusion which initiates cell migration has a significant amount of subcellular heterogeneity in micrometer length and minute time scales. It is driven by actin polymerization, e.g., pushing the plasma membrane forward, and then regulated by a multitude of actin regulators. While mathematical modeling is central to system-level understandings of cell protrusion, most of the modeling is based on the ensemble average of actin regulator dynamics at the cellular or population levels, preventing from capturing the heterogeneous cellular activities. With these in mind, a systematic modeling framework is proposed in this paper for predicting velocities of heterogeneous protrusion of migrating cells driven by multiple molecular mechanisms. The modeling framework is developed through the integration of the multiple AutoRegressive eXogenous (ARX) models employing probability density input variables. Unlike conventional ARX models, it provides an effective framework for modeling heterogeneous subcellular behaviors with complex nonlinearities and uncertainties of dynamic systems. To train and validate the proposed model, numerous subcellular time series are extracted from time-lapse movies of migrating PtK1 cells using spinning disk confocal microscope: The current edge velocities and fluorescent intensities of mDia1, actin at the leading edge are used as the input while the future cell edge velocities are selected as an output. It is demonstrated that the proposed approach is highly effective in predicting the future trends of heterogeneous cell protrusion. In particular, by capturing the various multiple activities from the dataset, it is expected that it would improve the understanding of the molecular mechanism underlying cellular and subcellular heterogeneity.

## 1. Introduction

Over the last decade, a mathematical modeling approach has been suggested for better understanding complex cellular systems (DiMilla et al. 1991). However, it is still very challenging to quantitatively predict system-level cellular behaviors because the relationships between the multiple inputs and multiple outputs are inherently nonlinear and time-varying (Karr et al. 2012). Moreover, cellular activities are often highly heterogeneous, meaning that cells exhibit multiple phenotypes in space and time and changing their phenotypes depending on their environment (Karr et al. 2012; Wang et al. 2018). Therefore, building precise mathematical models of heterogeneous cellular behaviors is very difficult since they can be driven by several distinct underlying mechanisms quantitatively or qualitatively, including (1) there are a number of uncertainties in measuring and processing input signals of subcellular activities: (2) it is challenging to derive a mathematic model of highly heterogeneous nonlinear dynamic systems where multiple modes of molecular mechanisms are included; and (3) the acquired datasets are incomplete and incoherent in general.

The time series model such as autoregressive (AR) models has been attracted a great attention in a variety of engineering fields. This approach has been successfully applied to various areas such as weather forecasting and financial market as well as cell biology (Jaqaman et al. 2006). This is because it allows researchers to model complex system behaviors without comprehensive information about the system due to a data-driven approach (Chon et al. 2001; Kim et al. 2016). However, the traditional AR models do not consider complex nonlinear dynamic problems with numerous uncertainties such as the multiple cellular or subcellular activities. Therefore, there is a limited ability to model heterogeneous cellular activities. With these in mind, a discrete fuzzy modeling (DFM) framework is proposed to predict the heterogeneous cell motions from the recruitment of actin assembly factors involved in the protrusion of the cell membrane. It can predict the future subcellular protrusion velocity using current protrusion velocity with uncertain inputs such as mDia1 and actin fluorescence intensity. It is created through the integration of multiple autoregressive exogenous input (ARX) models, fuzzy logic theory, data clustering schemes, and weighted least squares estimators, as shown in Figure 1: (1) the uncertainties of input variables are incorporated into the proposed model; (2) the heterogeneity of complex molecular interaction is considered in the proposed modeling framework; and (3) a well-established pre-processing is used to address the incomplete and incoherent measurements.

**Figure 1.**
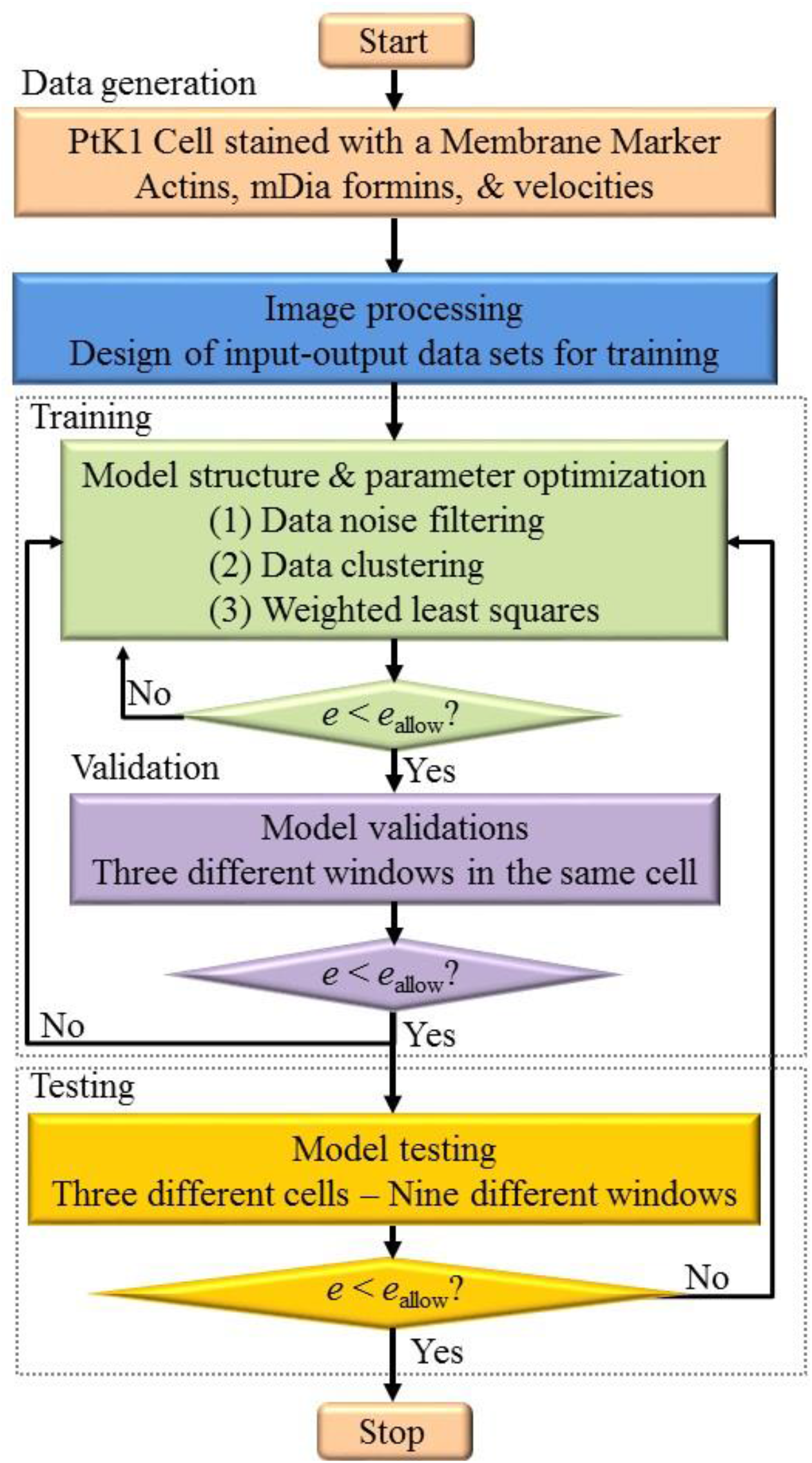
Proposed DFM algorithm.

## 2. Methods

### 2.1 Model architecture

The DFM model proposed in this paper is presented in Eq. (1) 
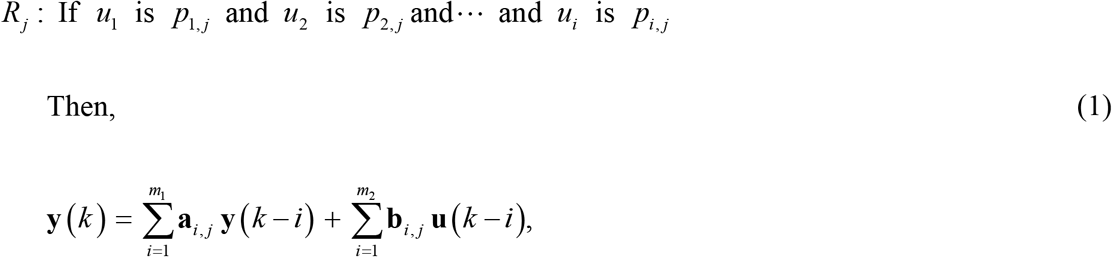
where *R_j_* is the *j^th^* rule and its consequent part is the *j^th^* linear dynamic model. A local dynamic system has a set of *if-then* rules. For example, “if the mDia1 is large, the velocities intensity increases”, or “if the actin is small, the velocities decrease.” *u_i_* is the *i^th^* premise variable, *p_i,j_* is the associated parameter, and *k* is the integer value. The output **y**(*k*)input **u**(*k*) and the associated parameter vectors **a**_*i*_ and **b**_*i*_ are presented in Eq. (2) to Eq. (5) 
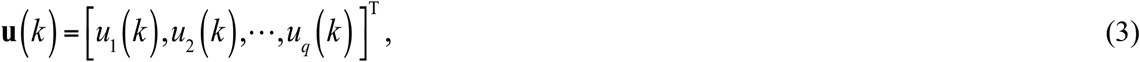

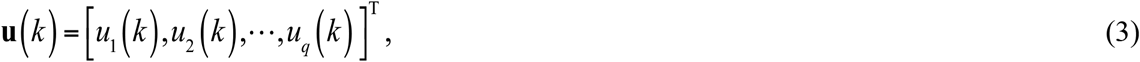

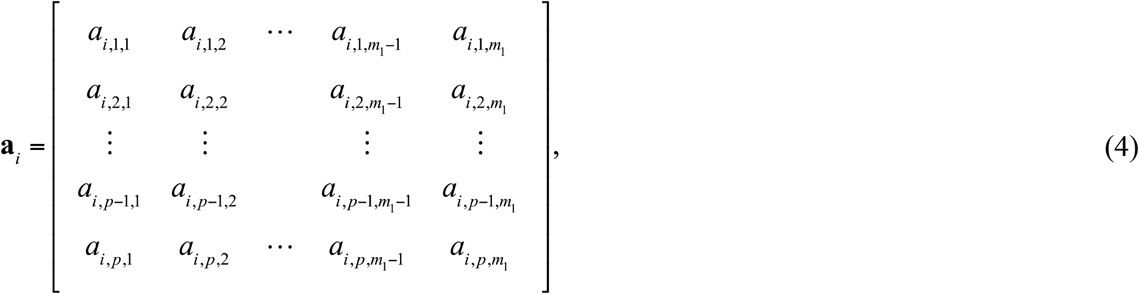

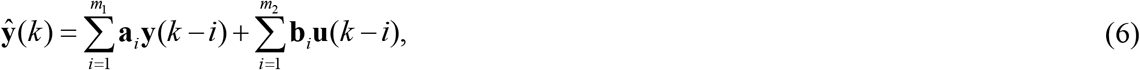
where *m*_1_ is the number of delay steps in the output; *m*_2_ is the number of delay steps in the input; **y**(*k*) is the output; **u**(*k*) is the input; *p* and *q* are the number of output and input variables, respectively, and **a***i* and **b***i* are the coefficient matrices to be estimated. In this study, *p*= 1 because the velocity is the only output signal. The multiple linear models at the specific operating point *u_i_* are integrated through a defuzzification approach (Kim *et al*. 2011; 2013; 2015; 2016). Therefore, Eq. (1) can be expressed as follows. 
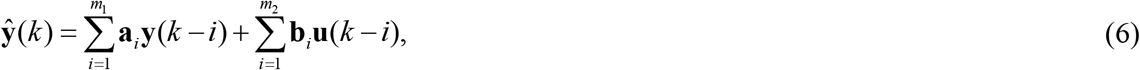
where 
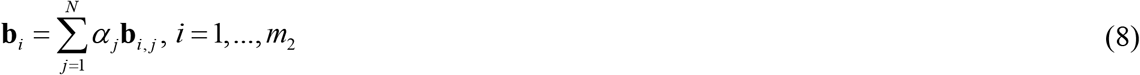

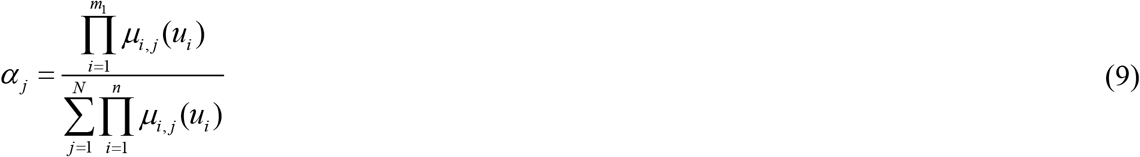
where 0 ≤ *α_j_* ≤ 1 is the normalized value of the *j^th^* rule and is expressed as shown in Eq. (9). 
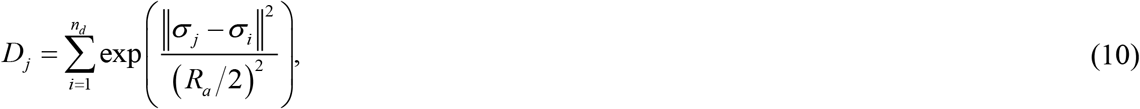
where *μ_i,j_* (*u_i_*) is the membership function (MF) of *u_i_* and *N* is the number of local dynamic models. MFs are useful to handle complex nonlinear systems with uncertain parameters. Fuzzy sets are constructed from the MFs. For example, if the level of mDia1 is categorized into three stages, *e.g*., low, medium, and high mDia1, a fuzzy set can be constructed. The MF for each antecedent variable in the DFM model should be carefully determined. In particular, the mean values of the probabilistic MFs need to be optimized. In this paper, a clustering algorithm is used.

### 2.2 Clustering algorithm

A cluster center is selected based on the highest density measure 
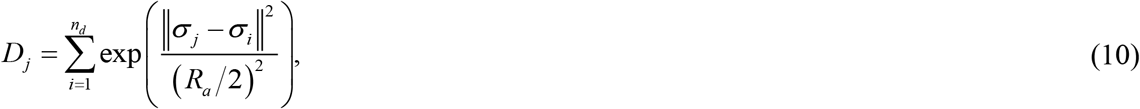
where *σ_i_* is the *i^th^* data point, *n_d_* is the total number of data points and *R_a_* is the range of data neighborhood. Subsequently, the selected cluster center and its neighborhood data points are reduced using the selection procedure 
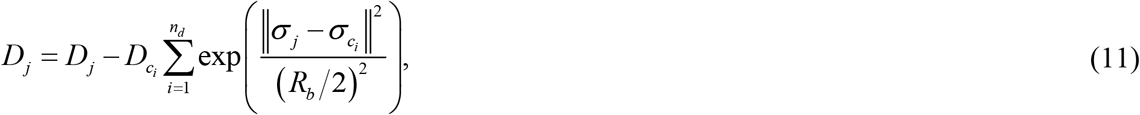
where *D_c_i__* is the *i^th^* density measure, *σ_c_i__* is the *i^th^* cluster center, and *R_b_* = *η_s_ R_a_* is used to avoid closely spaced centers where *η_s_* is a positive constant greater than 1. After subtraction, the cluster center is selected based on three criteria: (1) acceptance ratio, (2) rejection ratio, and (3) other relative distance criterion. This procedure is repeated until a sufficient number of cluster centers are found in the input space. The parameter settings for the subtractive clustering algorithm are assigned as *η_s_* is 1.25; *R_a_* is 0.35; the acceptance ratio is determined to be 0.5; and the rejection ratio is determined to be 0.15. It is noted that many different clustering algorithms would be available to estimate the ancedent parameters of the DFM (Babuska 1998; Lee et al. 2018). This clustering algorithm is used to construct the ancedent parameters. For example, the cluster center information is used as a center value of Gaussian or triangular MFs, as shown in Figure 2. Once the premise part of the DFM model is determined by the clustering algorithm, the weighted least squares algorithm is used to search optimum solutions of the consequent part parameters of the DFM model.

**Figure 2.**
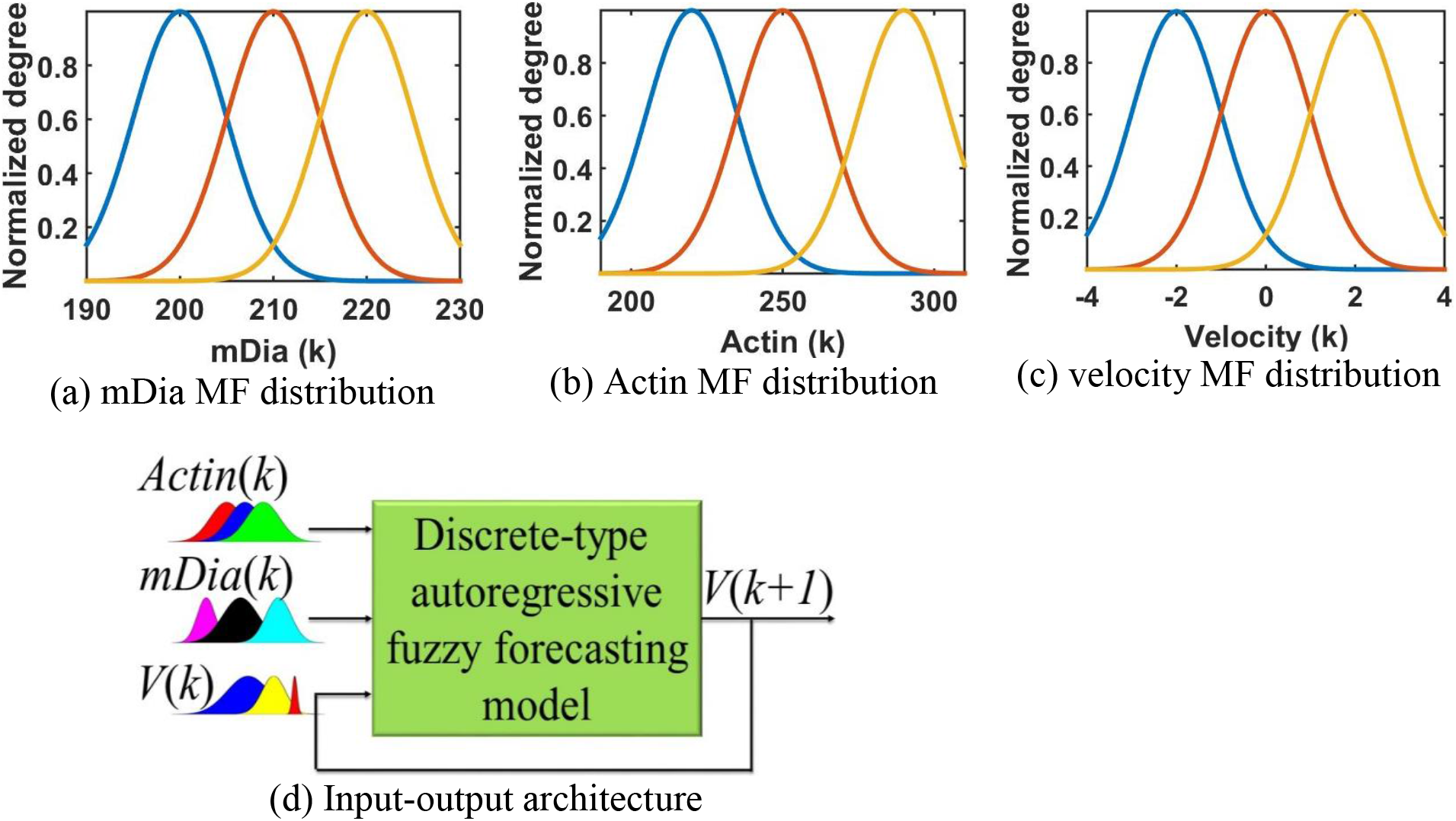
Configuration of the proposed DFM system. (a) Probabilistic description of the current mDia expressed in terms of fuzzy MF. (b) Probabilistic distribution of the current actin. (c) Probabilistic distribution of the current cell motion velocity. (d) An input-output system structure employing an output feedback.

### 2.3 Weighted Least Squares

The consequent part parameters of the DFM model are determined using the weighted linear least squares 
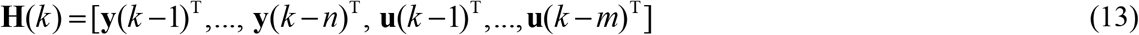
where 
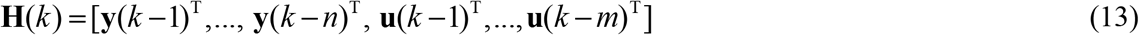

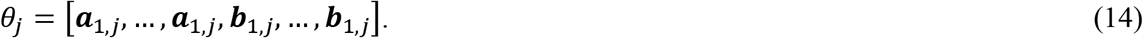
and 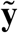(*k*) is the measured data, *w*_*j*_ is the weighting factor.

### 2.4 Proposed DFM algorithm

The DFM algorithm proposed in this paper is as follows. The flowchart for the proposed algorithm is depicted in Figure 1.

Step 1: The images on dynamic motions are collected, and then various components such as the mDia1, actins, and velocities are extracted from the collected images.

Step 2: Correlation coefficients between each pair of input and output signals are calculated. The signals with high coefficient values are retained as input signals. To reduce the number of input signals, partial correlation coefficients are calculated to determine which signals can be removed from the data sets.

Step 3: The clustering algorithm is used to construct the premise part of the DFM model.

Step 4: Once the antecedent part are determined using the clustering algorithm, the consequent parameters are optimized using the weighted least squares algorithm.

Step 5: The performance of the DFM model is estimated via various evaluation indices, including percent error in peak, bias, mean square error, root mean square error, and coefficient of determination. If the modeling performance is not satisfied (*i.e*. the errors are larger than the allowable error limits), the modeling process goes to Step 3. Note that Step 3 to Step 5 are repeated until the errors converge to desirable values. For example, the number of membership functions can be adjusted. The target errors are determined by users.

Step 6: When the model estimates are satisfied with the specified boundaries of errors, the model is tested using other data sets that are not used for the training process. In this paper, the specified boundaries of errors are determined qualitatively by visual inspection of the time series as well as quantitatively by the evaluation index such as *R*^2^. If the prediction is not satisfied, the procedure goes to Step 3. If it is satisfied, the algorithm would stop.

Note that trial and error is required for Step 3 to Step 6. When it is difficult to develop an effective model from Step 3 to Step 6, it is recommended to return to Step 2. Based on different combinations of input-output signals, it is sometimes possible to improve the modeling performance. It should be noted that the computational costs of calculating the output significantly increase when the number of input variables grows, making the DFM model in high dimension unfeasible. It is often counterproductive to consider a high number of input variables in the prediction model for a restricted purpose.

### 2.5 Experimental data

Sample videos for the analysis were prepared by taking time-lapse movies of PtK1 cells expressing fluorescently tagged mDia1 and actin with a spinning disk confocal microscope for approximately 200 frames at 5 sec/frame, as shown in Figure 3. After segmenting the leading edge of each cell by multiple probing windows with an area of 1 μm^2^, time series of velocities and fluorescence intensities of the tagged mDia1 and actin acquired from each probing window were quantified (Machacek *et al*. 2009; Lee *et al*. 2015). These time series datasets are used for the DFM.

**Figure 3.**
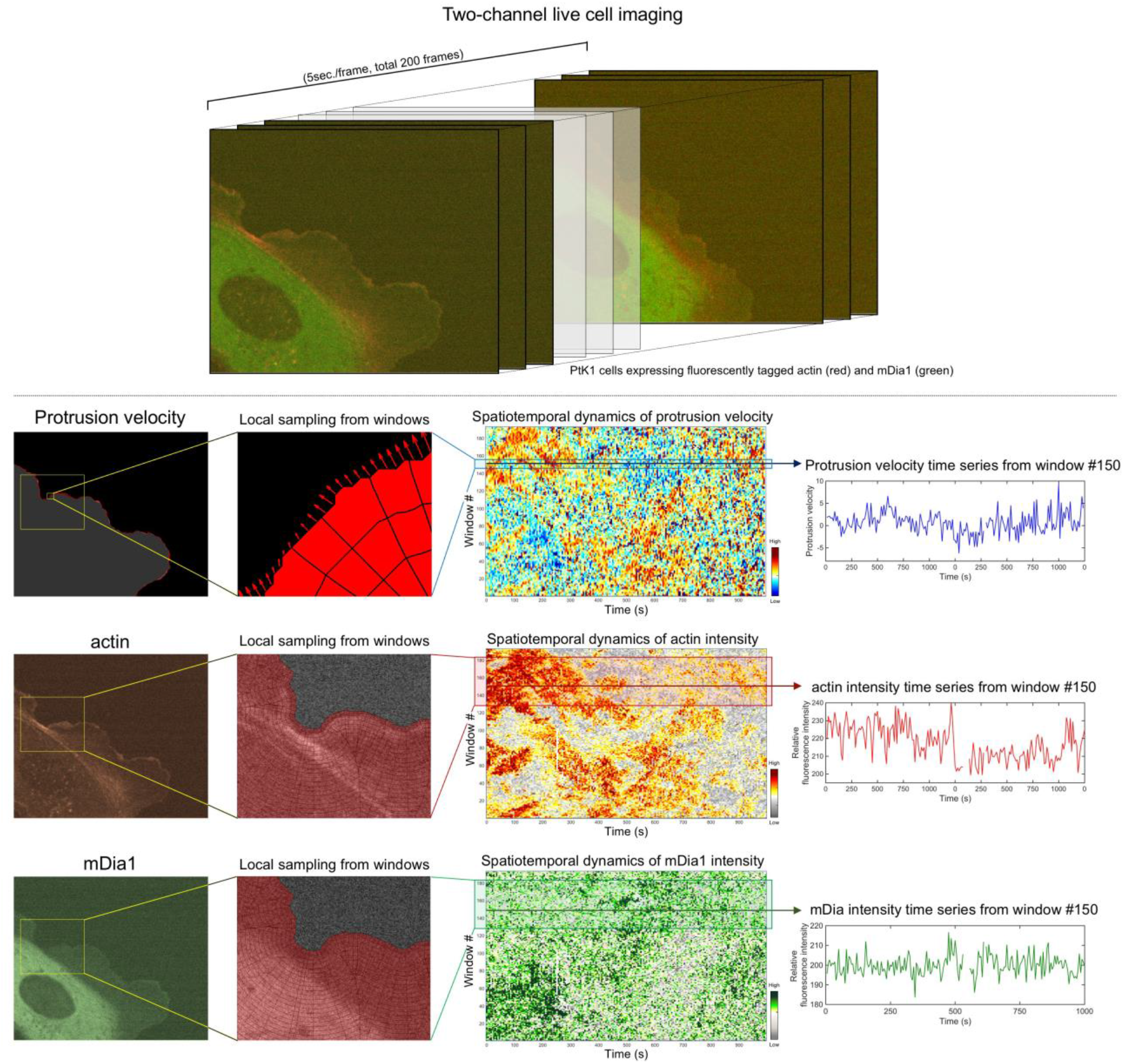
A local sampling of protrusion velocities and fluorescence intensities of actin and mDia1 at the leading edge of migrating PtK1 Epithelial cells.

The dataset used in this study contains a significant amount of heterogeneity. First, recently we identified six different protrusion phenotypes by deconvolving the subcellular protrusion heterogeneity using unsupervised learning (Wang *et al*, 2018). Moreover, the leading edge of the cells undergoes protrusion and retraction cycles. The protrusion and retraction are distinct processes, which are driven by the different molecular mechanism. For example, the protrusion is driven by actin assembly processes whereas the retraction is driven by myosin contraction. In our DFM framework, the model training was performed without knowing that the time series are in the protrusion or retraction phases. Second, there are at least five different protrusion phenotypes based on our previous clustering analyses of the protrusion velocities (Wang *et al*, 2018).

## 3. RESULTS

The input of the DFM includes the fluorescence intensity of mDia1 and actin, and current velocities while the future velocities are used as an output signal. To further evaluate the effectiveness of the trained DFM model, a variety of different validation datasets that are not used in the training process are applied to the trained model. In this study, both qualitative (i.e., visual inspections) and quantitative analysis methods are used to evaluate the performance of the proposed modeling framework: the modeling errors are visually present first and then they are quantified using several indices.

### 3.1 Qualitative analysis of numerical model

The performance of the DFM models can be judged first by visual inspection (i.e., viewing patterns in data). It is easily to detect under- or non-modeled patterns and capture the overall behavior of the model without conducting the extensive quantitative analysis. In many problems, simple visual inspection of models is sufficient (Bennett et al. 2013; Kim et al. 2015). In this study, the time series prediction, the residual, quantile-quantile (QQ), and normal probability density function plots are used. Figure 4 (a) and (b) compare the predicted velocities (Model) with the measured training and validated datasets, respectively. DATAt represents the measured data used for the model training while DATAv is the validation data. The solid red line is the model while the dotted black lines are datasets. As shown in the figure, great agreements between the predicted values and measurements are found. Figure 4 (c) and (d) shows the residual error plots for the trained model and its validated results. The residual errors of both training and validation models appear random, which suggests that there are no systematic errors in the models. For instance, high density of positive/negative values is not found in the plots, which indicates that all the models do not tend to over/under-estimate the measured values (Bennett et al. 2013). Figure 4 (e) presents the QQ plots of the trained model and measured data while Figure 4 (f) is the QQ plot of the proposed model and validation data. If the model and data come from the same distribution, the QQ plot will be linear. From Figure 4 (e) and Figure 4 (f), all the QQ plots are closely linear, which means that both models and datasets come from the same distribution. These QQ plots correspond to the normal distribution functions (NDF) in Figure 4 (g) and (h). As shown in the figure, good agreements between the model NDF and the measured data are found for both the training and validation. The error analysis is quantitatively conducted in next section.

**Figure 4.**
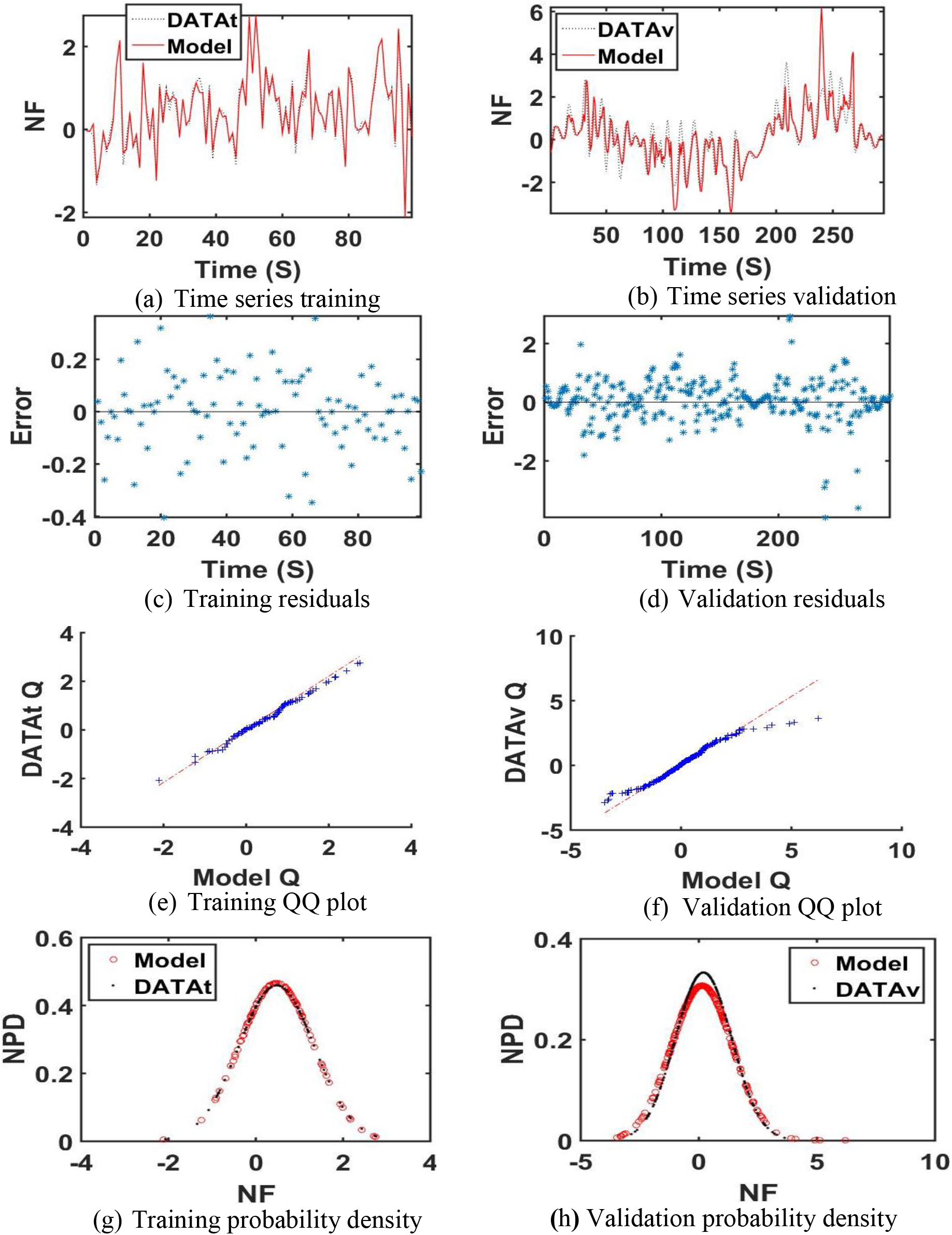
Visual inspection of the proposed model. (a): the prediction velocities of the trained DFM. (b): the validated DFM using different cell datasets. (c): the residual error for the trained DFM model. Figure 4 (d): the residual error for the validated results. (e): the quantile-quantile (QQ) plot of the trained DFM model. (f): the QQ plot of the validated data. (g): the comparison of normal distribution functions (NDF) of the DFM and training data. (h): the NDF comparison of the DFM and validation data.

### 3.2 Quantitative analysis of numerical model

In order to quantify the modeling error, several evaluation indices were used. The simulation results are shown in Table 1. As shown in Table 1, the proposed DFM model is effective in predicting the complex behavior of cell motion fluctuations. The trained model demonstrates good performance according to all the indices. The maximum errors of the trained model in peak (*J*_1_) in forecasting cell motions are smallest compared to the other validation models. The *J*_1_ metric in the validation process is negative because the DFM model slightly overestimates the overall data values by 0.17%. The validation error is slightly higher than the training error, as measured by *J*_1_.

**Table 1.** Quantitative evaluation of the proposed model.

| Data Type | $J_1$ | $J_2$ | $J_3$ | $J_4$ | $J_5$ |
| --- | --- | --- | --- | --- | --- |
| TRAINING | 0.000231064 | 1.48446e-06 | 0.021043 | 0.145062 | 0.971827 |
| Validation 01 | -0.742045 | 0.0077347 | 0.414318 | 0.643676 | 0.723545 |
| Validation 02 | -0.228814 | -0.0100591 | 0.300402 | 0.548089 | 0.724686 |
| Validation 03 | 0.0357843 | -0.00152822 | 0.316371 | 0.562469 | 0.72075 |
| Validation 04 | -0.0140609 | -0.0231869 | 0.274078 | 0.523525 | 0.712479 |
| Validation 05 | 0.29496 | -0.037906 | 0.356235 | 0.596854 | 0.730172 |
| Validation 06 | -0.179856 | 0.0347253 | 0.197477 | 0.444384 | 0.733802 |
| Validation 07 | -0.231245 | -0.0425631 | 0.371844 | 0.60979 | 0.707997 |
| Validation 08 | -0.0693903 | -0.054403 | 0.387021 | 0.62211 | 0.726629 |
| Validation 09 | -0.186876 | 0.0121289 | 0.262823 | 0.512663 | 0.702281 |

**Note:** $\hat{y}$ : forecasting values, $\tilde{y}$ : the data measured using the microscopy, $\bar{y}$ : the mean value of the measured data. $N_t$ : the number of data points. $J_1 = \left[ \max(\tilde{y}_i) - \max(\hat{y}_i) \right] / \max(\tilde{y}_i) \times 100$ : determine if the DFM models can produce a similar range of observed data. $J_2 = \sum_{i=1}^{N_t} (\tilde{y}_i - \hat{y}_i) / N_t$ : the mean of the residuals as the bias. $J_3 = \sum_{i=1}^{N_t} (\tilde{y}_i - \hat{y}_i)^2 / N_t$ : mean square errors adopted to determine if the DFM models tend to overestimate and underestimate the measured data. $J_4 = \sqrt{\sum_{i=1}^{N_t} (\tilde{y}_i - \hat{y}_i)^2 / N_t}$ : the root mean square error to express the normalized error metric. $J_5 = 1 - \left[ \sum_{i=1}^{N_t} (\tilde{y}_i - \hat{y}_i)^2 / N_t \right] / \left[ \sum_{i=1}^{N_t} (\tilde{y}_i - \bar{y})^2 / N_t \right]$ : the coefficient of determination to see how well the DFM models capture the variance in the measured data.

This is because the highest peak error in the validated model is higher than the one in the training time history data. As seen in *J*_2_, the trained model slightly overestimates some validated data. However, the occurrence of both positive and negative errors in *J*_2_ could result in a value close to zero, thus indices *J*_3_ and *J*_4_ account for this issue. As seen in *J*_3_, the performances of all the trained models are better than the validation models. The RMSE provides a normalized metric, yielding values between 0.14 and 0.64, for all models, respectively (*J*_4_). The coefficient of determination (*J*_5_) for the proposed DFM models is 97% for the training data, indicating strong agreement. The coefficients of determination of the DFM models range from 70% to 73% for the validation data.

## 4. Discussion

In this paper, a time series model is proposed for predicting the subcellular heterogeneous protrusive motion of migrating cells. The prediction model is developed through the integration of multiple autoregressive models, fuzzy logic membership functions, data clustering algorithms, and multiple weighted least squares algorithms. The discrete-type fuzzy model (DFM) was trained using the actin, mDia1, and current velocities as input signals and the future velocities as an output. The trained model was validated using different datasets collected from 9 different locations in the same cell. It is demonstrated from extensive experiments (both experimental and numerical testing) that the proposed DFM is very effective in predicting the complex nonlinear behavior of cellular systems.

It was observed that it is effective in modeling subcellular protrusion heterogeneity when mDia1, actin, and edge velocity are used as training datasets. It is believed that mDia1 initiates the protrusion by nucleating actin filaments that other actin nucleators and Arp2/3 complex can bind (Isogai *et al*. 2015; Lee *et al*. 2015). Since that mDia1 is not a major driver of actin polymerization for cell protrusion, it can be inferred from this fact that mDia1 may be an important player which generates the protrusion heterogeneity.

It is highly expected that the proposed modeling framework can improve the mechanistic understanding of heterogeneous cellular and subcellular behaviors, extracted from live cell imaging data, and can be applied to the other cellular and subcellular heterogeneity in cellular processes such as cell migration, cell division, cytoskeletal structures, and membrane-bound organelles.

## Acknowledgments

This work was supported by the NIH (Grant Number: GM122012), CBU, and WPI.

## Author Contributions

Y.K and K.L. conceived and initiated the project. Y. K. implemented the DFM algorithm into the subcellular datasets. H. C. designed, conducted the experiments, and validated the results. Y. K. and K. L. coordinated the study and wrote the final version of the manuscript and supplement. All authors discussed the results of the study.

## References

Arsava, S.K., Kim, Y., El-Korchi, T. and Park, H.S. (2013) Nonlinear system identification of smart structures under high impact loads, Journal of Smart Materials and Structures, 22, doi:10.1088/0964-1726/22/5/055008.

Arsava, S.K., Chong, J.W. and Kim, Y. (2014) A novel health monitoring scheme for smart structures, Journal of Vibration and Control, doi: 10.1177/1077546314533716.

Arsava, S.K., and Kim, Y. (2015a) Modeling of magnetorheological dampers under various impact forces, Shock and Vibration, vol. 2015, Article ID 905186, 20 pages, 2015. doi:10.1155/2015/905186

Arsava, S.K., Nam, Y. and Kim, Y. (2015b) Nonlinear system identification of smart reinforced concrete structures under high impact loads, Journal of Vibration and Control, doi: 10.1177/1077546314563966

Arsava, S.K., Kim, Y., Kim, K.H. and Shin, B.S. (2015c) Smart fuzzy control of reinforced concrete structures excited by collision-type forces, Expert Systems with Applications, 42(21), 7929–7941.

Cha, Y.J., Agrawal, A.K., Kim, Y. and Raich, A. (2012) Multi-objective genetic algorithms for cost-effective distributions of actuators and sensors in large structures, Expert Systems with Applications, 39, 7822–7833.

Cha, Y.J., Kim, Y., Raich, A. and Agrawal, A.K. (2013) Multi-objective optimization for actuator and sensor layouts of actively controlled 3D buildings, Journal of Vibration and Control, 19, 942–960.

Chong, J.W., Kim, Y. and Chon, K. (2014) Nonlinear multiclass support vector machine-based health monitoring system for buildings employing magnetorheological dampers, Journal of Intelligent Material Systems and Structures, 25, 1456–1468

DiMilla, P.A., Barbee, K., and Lauffenburger D.A. (1991) Mathematical model for the effects of adhesion and mechanics on cell migration speed, Biophysical Journal, 60, 15–37.

Hughes, J.E., Kim, Y. and El-Korchi, T. (2015) Radar technology for structural hazard mitiga-tion, Journal of Vibration and Control, 2015, In Press.

Karr, J.R., Sanghvi, J.C., Macklin, D.N., Gutschow, M.V., Jacobs, J.M, Bolival B., Assad-Garcia, N., Glass, J.I., and Covert, M.W., (2012) A Whole-Cell Computational Model Predicts Phenotype from Genotype, Cell 150, 389–401.

Isogai, T., van der Kammen, R., Leyton-Puig, D., Kedziora, K. M., Jalink, K., & Innocenti, M. (2015). Initiation of lamellipodia and ruffles involves cooperation between mDia1 and the Arp2/3 complex. J Cell Sci, jcs-176768.

Kim Y. and Langari R. (2007) Nonlinear Identification and Control of a Building Structure with a Magnetorheological Damper System. American Control Conference, New York, July 11–13.

Kim Y., Langari R. and Hurlebaus S. (2009b). Semiactive Nonlinear Control of a Building Using a Magnetorheological Damper System. Mechanical Systems and Signal Processing 23, 300–315.

Kim Y., Hurlebaus, S. and Langari, R. (2010a) Control of a seismically excited benchmark building using linear matrix inequality-based semiactive nonlinear fuzzy control,¡± ASCE Journal of Structural Engineering, 136(8), 1023–1026.

Kim Y., Langari, R., and Hurlebaus, S. (2010b) Model-based multi-input, multi-output supervisory semiactive nonlinear fuzzy controller, Computer-Aided Civil and Infrastructure Engineering, 25, 387–393.

Kim Y., Kim, C. and Langari, R. (2010c) Novel bio-inspired smart control for hazard mitigation of civil structures, Journal of Smart Materials and Structures, 19, 115009, doi:10.1088/0964-1726/19/11/115009.

Kim Y., Hurlebaus S., Sharifi R. and Langari R. (2009a) Nonlinear Identification of MIMO Smart Structures ASME Dynamic Systems and Control Conference Hollywood, California, Oct. 12–14.

Kim Y., Hurlebaus S, and Langari, R. (2011) Fuzzy Identification of Building-MR Damper System International Journal of Intelligent and Fuzzy Systems, 22(4), 185–205.

Kim Y., Chong, J.W., Chon, K. and Kim, J.M. (2013) Wavelet-based AR-SVM for health monitoring of smart structures, Journal of Smart Materials and Structures, 22(1), 015003, doi:10.1088/0964-1726/22/1/015003.

Kim Y., Mallick, R., Bhowmick, S. and Chen, B. (2013) Nonlinear system identification of large-scale smart pavement systems,¡± Expert Systems with Applications, 40, 3551–3560.

Kim Y., Kim, K.H. and Shin, B.S. (2014a) Fuzzy model forecasting of offshore bar-shape profiles under high waves,¡± Expert Systems with Applications, 41, 5771–5779.

Kim Y., Bai, J.W. and Albano, L.D. (2014b) Fragility estimates of smart structures with sensor faults, Journal of Smart Materials and Structures, 22 125012, doi:10.1088/0964-1726/22/12/125012.

Kim Y., Shin, S.S. and Plummer, J.D. (2014c) A wavelet-based autoregressive fuzzy model for forecasting algal blooms,¡± Environmental Modeling & Software, 62, 1–10

Kim Y., Kim, Y.H. and Lee, S. (2015) Multivariable nonlinear identification of smart buildings, Mechanical Systems and Signal Processing, 62–63, 254–271.

Lee K, Elliott, H.L., Oak, Y., Groisman, A., Tytell, J.D., and Danuser, G. (2015) Functional hierarchy of redundant actin assembly factors revealed by fine-grained registration of intrinsic image fluctuations, Cell Systems, 1, 37–50, http://dx.doi.org/10.1016/j.cels.2015.07.001

Machacek, M., Hodgson, L., Welch, C., Elliott, H., Pertz, O., Nalbant, P., … & Danuser, G. (2009). Coordination of Rho GTPase activities during cell protrusion. Nature, 461(7260), 99.

Mitchell R., Kim, Y., El-Korchi, T. and Cha, Y.J. (2013) Wavelet-neuro-fuzzy control of hybrid building-active tuned mass damper system under seismic excitations, Journal of Vibration and Control, 19(12), 1881–1894

Mitchell R., Kim, Y., and El-Korchi, T. (2012) System identification of smart structures using a wavelet neuro-fuzzy model, Journal of Smart Materials and Structures, 21, doi:10.1088/0964-1726/21/11/115009.

Mitchell R., Cha, Y.J., Kim, Y. and Mahajan, A. (2015) Active control of highway bridges subject to a variety of earthquake loads, Earthquake Engineering and Engineering Vibration, 14(2), 253–263.

Mohammadzadeh, S., Kim, Y. and Ahn, J. (2015) PCA-based neuro-fuzzy model for system identification of smart structures, Journal of Smart Structures and Systems, 15(4), 1139–1158.

Sharifi, R., Kim, Y. and Langari, R. (2010) Sensor fault isolation and detection of smart structures Journal of Smart Materials and Structures 19 doi:10.1088/0964-1726/19/10/105001

Wang, C., Choi, H. J., Kim, S. J., Desai, A., Lee, N., Kim, D., Bae Y., and Lee, K. (2018) Deconvolution of subcellular protrusion heterogeneity and the underlying actin regulator dynamics from live cell imaging. Nature Communications, 9. 1688.

